## Supplementary Figures for "Heterogeneous somatostatin-expressing neuron population in mouse ventral tegmental area"

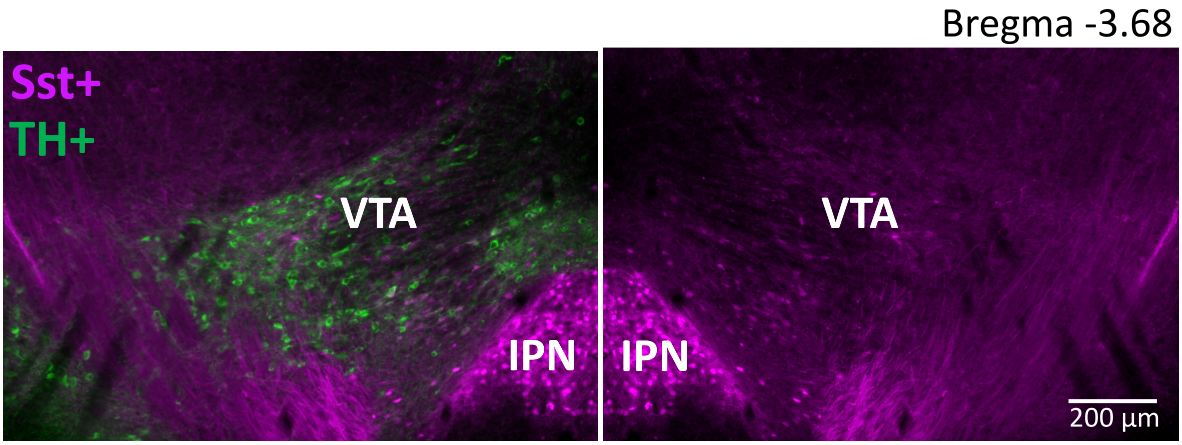


**Figure S1. Anatomical localization of Sst neurons within the VTA in coronal plane**. A representative image of coronal sections, which were used for IHC-based counting. The VTA was defined by TH+ staining (excluding the substantia nigra compacta) – left image. Right image shows the magenta Sst cells in the same section (horizontally flipped) with the green TH+ channel off.

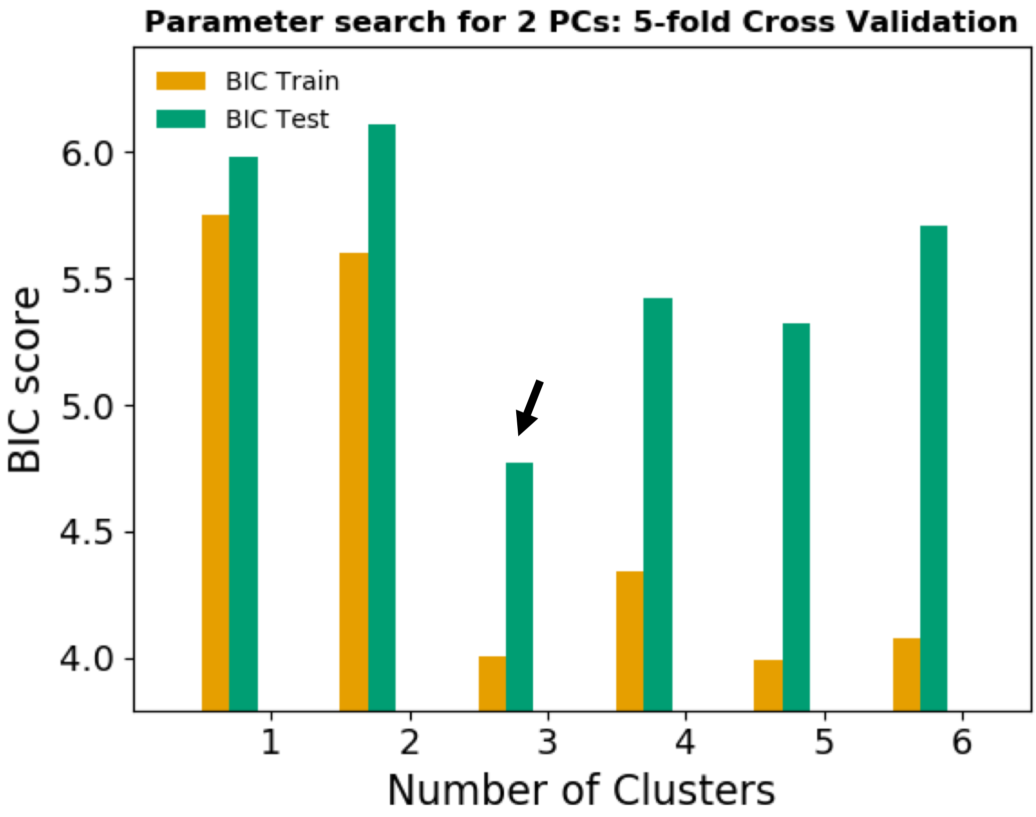


**Figure S2. Bayesian information criterion (BIC) for model selection**. According to the BIC, the combination of 2 principal components (PCs) and 3 clusters gives the best fit to describe the data (black arrow). Orange bars represent the BIC score for the model training dataset; green bars represent the BIC score for the test dataset unfamiliar to the model.

a

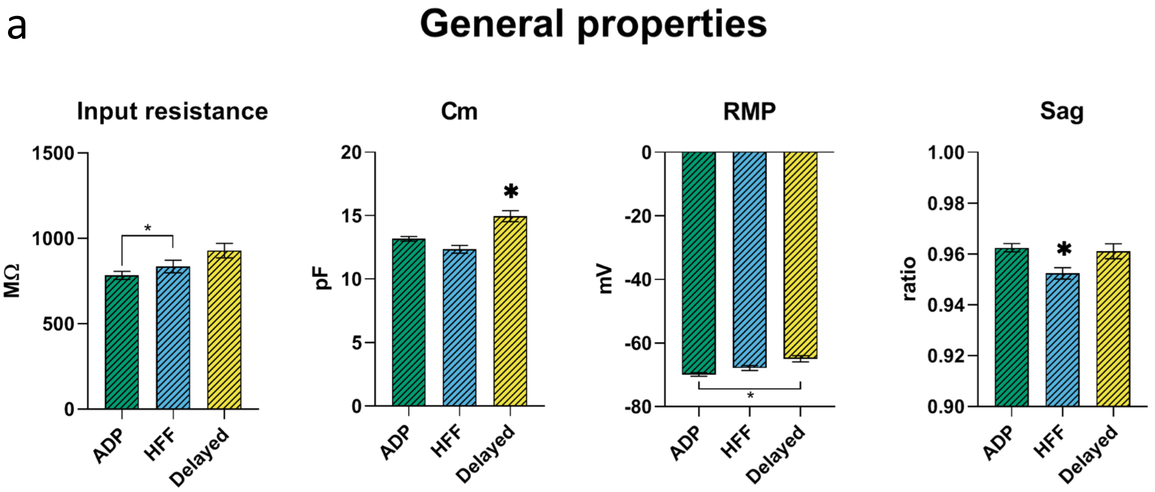


**Figure S3. Passive and active electrophysiological properties of VTA Sst-neuron subtypes.**

**a.** General membrane properties. Cm, cell capacitance; RMP, resting membrane potential. **b.** Electrophysiological properties of the first action potential (AP) at rheobase level of excitation. ADP, afterdepolarization; AHP; afterhyperpolarization; HW, half-width. **c.** Electrophysiological properties of the subtypes at the saturating level of excitation, producing the highest number of action potentials. All graphs show means ± SEM for ADP (n=215), HFF (n=92) and Delayed (n=85) neurons. Statistical significances between groups were measured by one-way ANOVA with Tukey's post hoc test. Big asterisks show values, which are significantly different from two others ($p < 0.05$). Small asterisks with the connecting line indicate only two significantly different values.

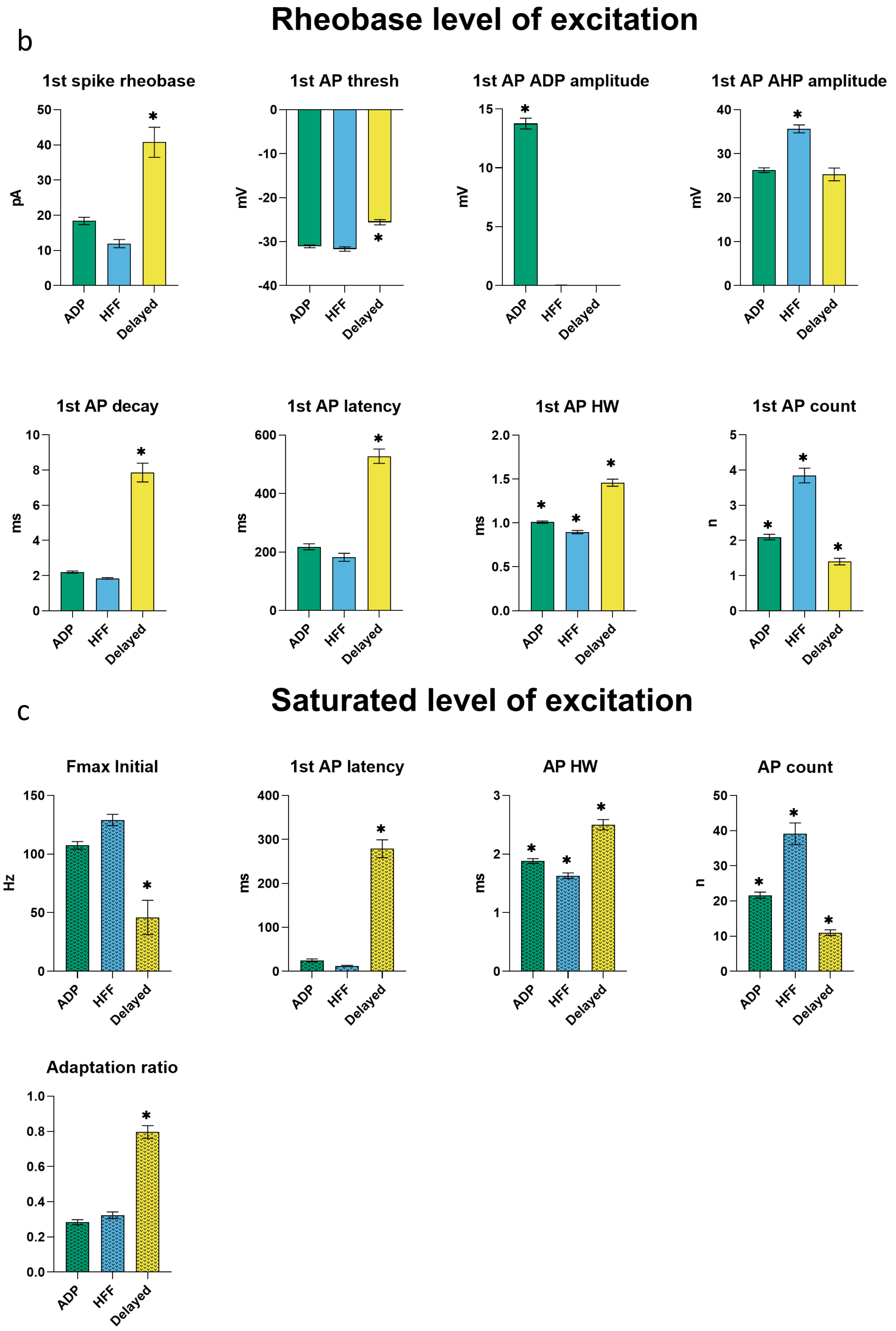

Rheobase level of excitation
b
1st spike rheobase
pA
1st AP thresh
mV
1st AP ADP amplitude
mV
1st AP AHP amplitude
mV
1st AP decay
ms
1st AP latency
ms
1st AP HW
ms
1st AP count
n
ADP
HFF
Delayed
Saturated level of excitation
c
Fmax Initial
Hz
1st AP latency
ms
AP HW
ms
AP count
n
Adaptation ratio
ADP
HFF
Delayed

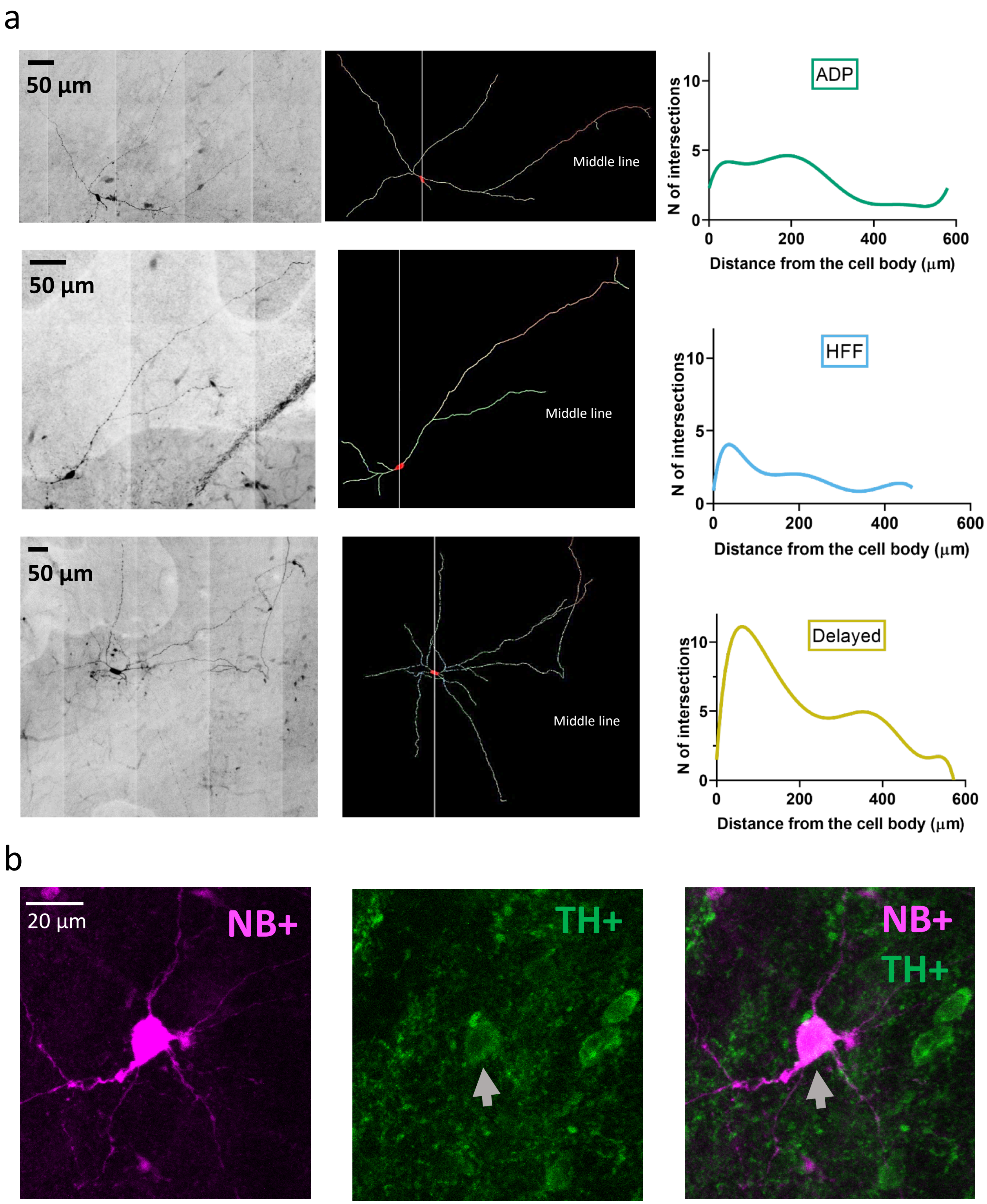


**Figure S4. Examples of VTA Sst-neuron morphology. a.** From the left, line by line: black and white-inverted copies of original micrographs of neurobiotin-filled Sst cells of the ADP, HFF and Delayed subtypes defined by clustering according to their electrophysiological features. Middle: reconstructed morphology of the same neurons. Right: the Sholl curves for the same cells. **b.** Example of a neurobiotin-filled (NB+) Delayed neuron with positive TH staining.

a

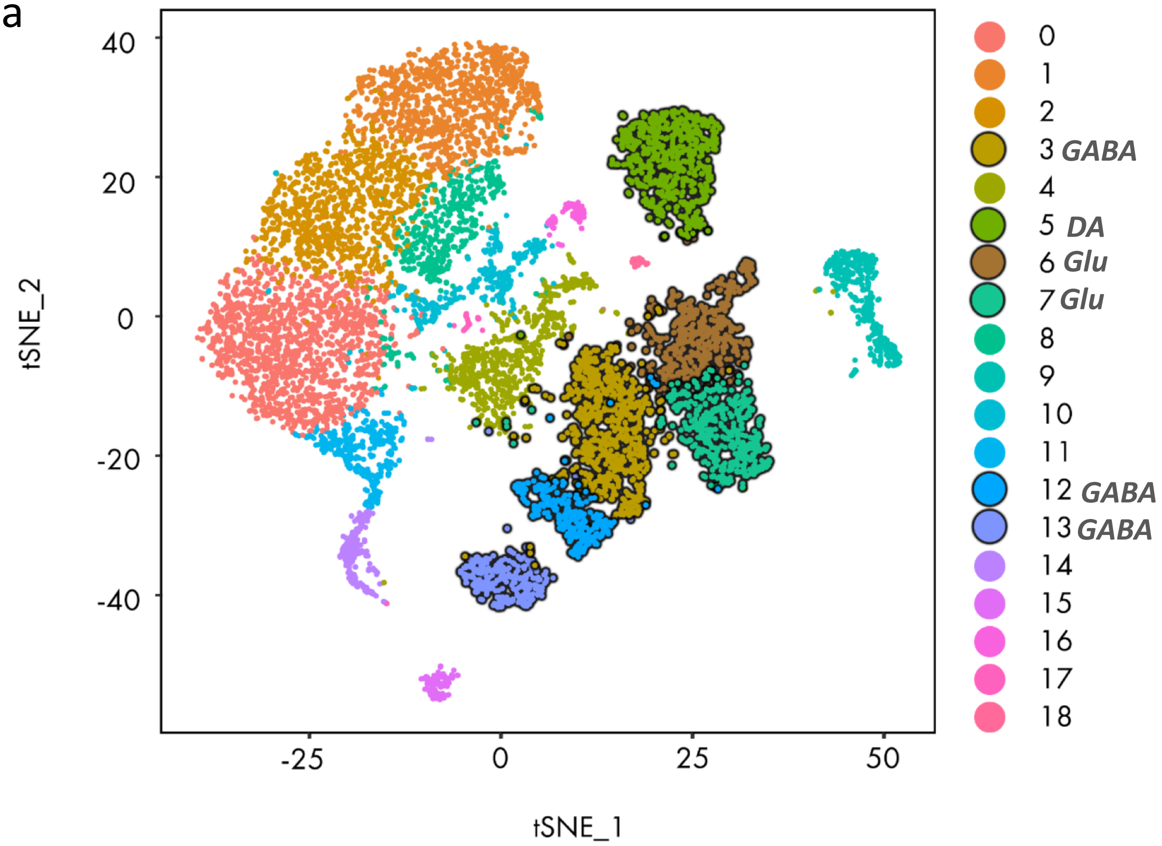

40
20
0
-20
-40
tSNE_2
-25
0
25
50
tSNE_1
0
1
2
3 GABA
4
5 DA
6 Glu
7 Glu
8
9
10
11
12 GABA
13 GABA
14
15
16
17
18


b

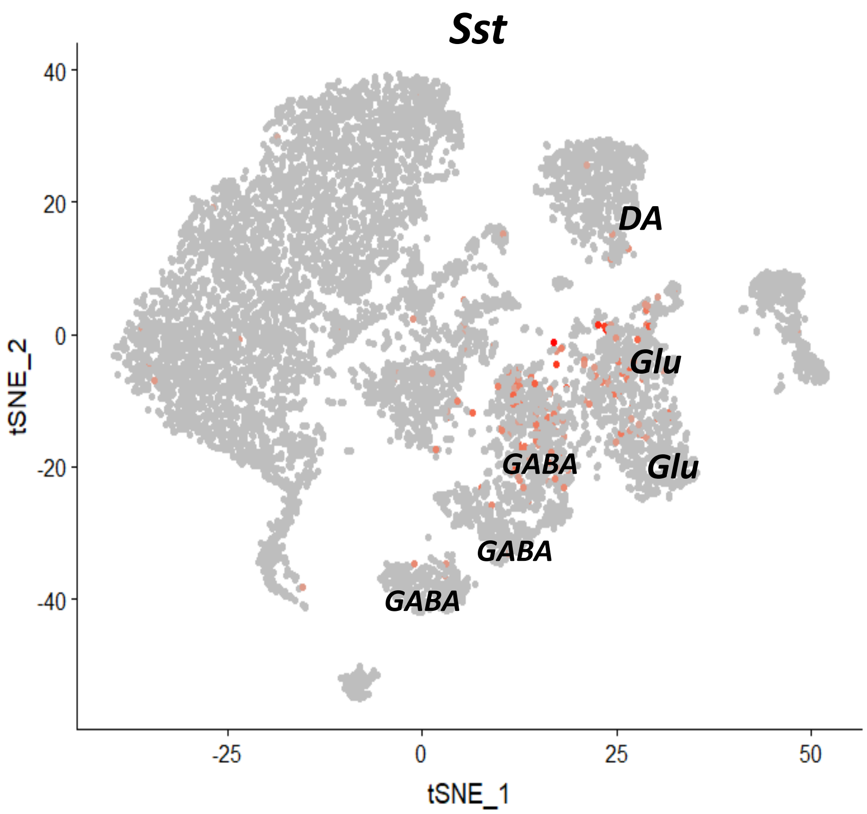

Sst
40
20
0
-20
-40
tSNE_2
DA
Glu
GABA
Glu
GABA
GABA
-25
0
25
50
tSNE_1

c

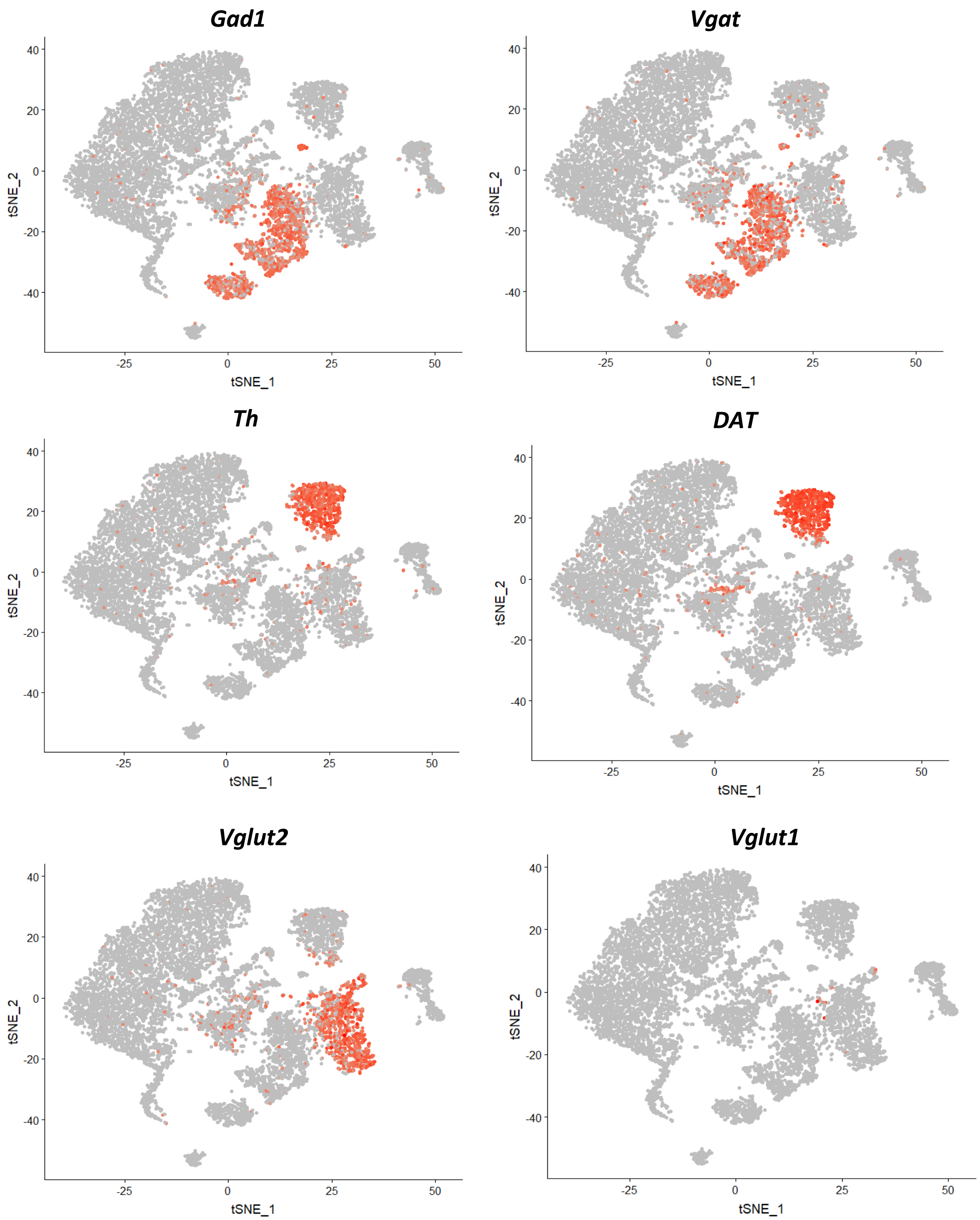


**Figure S5. Seurat clustering of the reference midbrain dataset and selection of the clusters for PatchSeq cells classification. a.** tSNE of the clustering results. Highlighted with dark contour clusters were used for the mapping procedure. Clusters 3, 5, 6, 7, 12 and 13 were selected due to Sst (panel **b**) and neuronal markers expression (panel **c**).

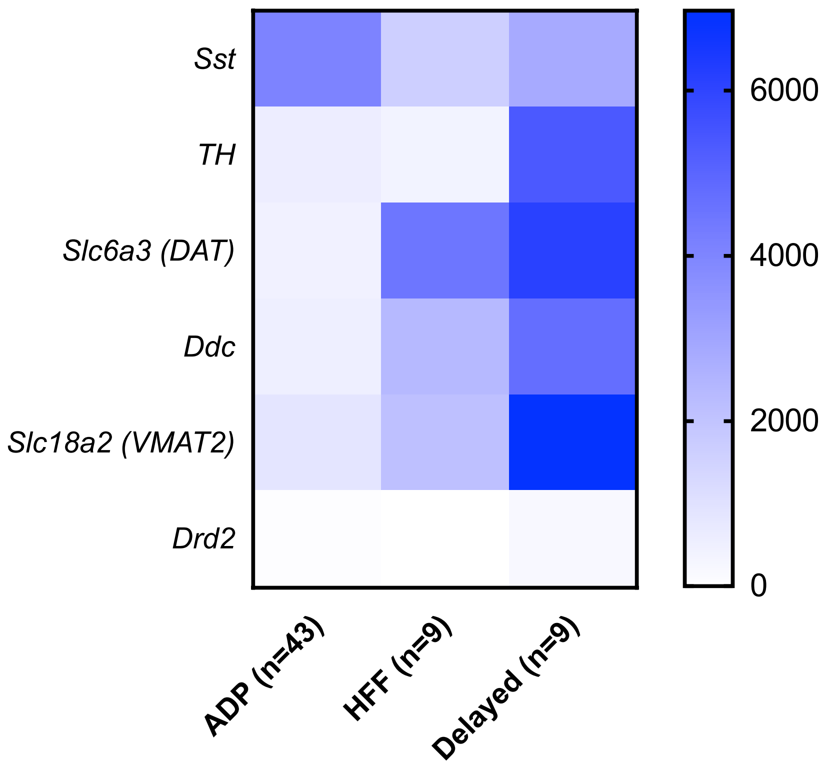


**Figure S6. Expression of dopamine-related genes in different Sst-expressing neuronal populations in the VTA, indicating strong expression in Delayed cells.** Values are mean expression levels in read counts for n cells. For scaling, the values for *TH* were divided by 10.

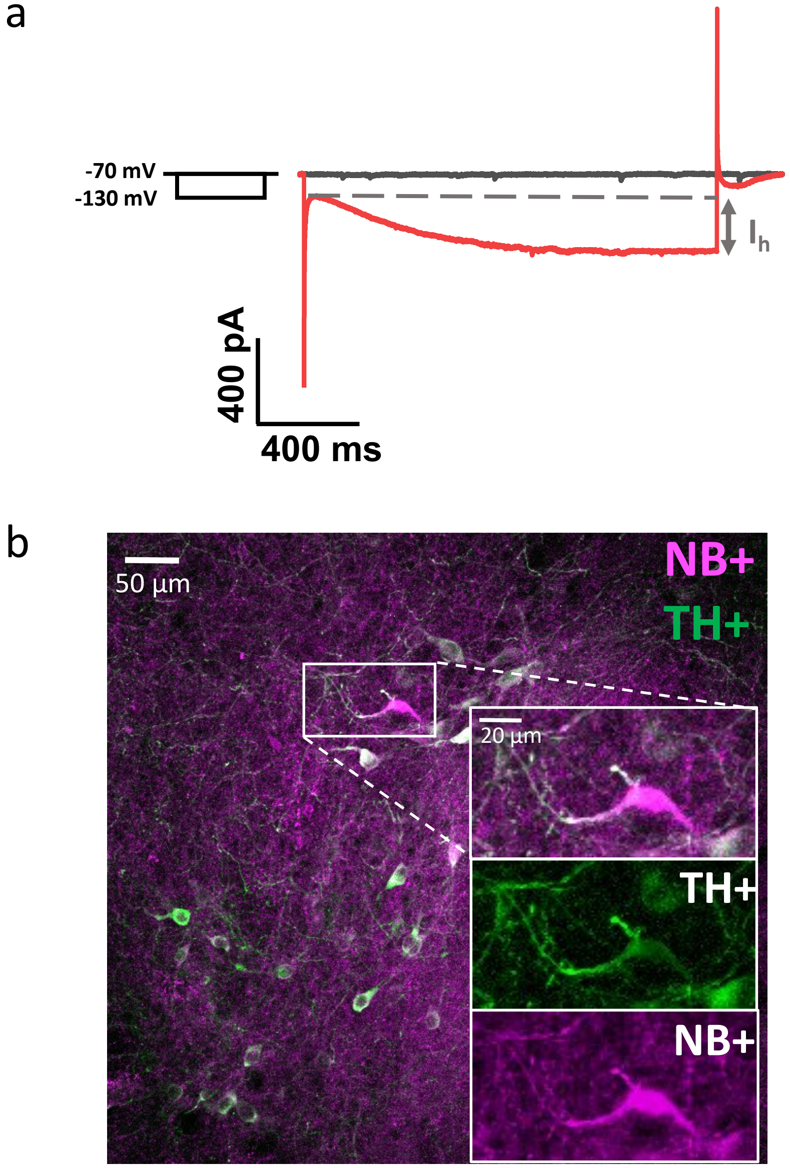


**Figure S7. Examples of $I_h$-current test (a) and immunohistochemistry (b) for confirmation of DA neuron phenotype. a.** $I_h$ current is shown as an extra inward current, occurring during hyperpolarization of the cell membrane potential from -70 to -130 mV. **b.** The recorded DA neuron was positive for Neurobiotin (magenta) and TH (green). Often TH was leaking out from the cell body during neurobiotin loading, explaining why many cells are brighter co-stained at axonic/dendritic sites than at somas.

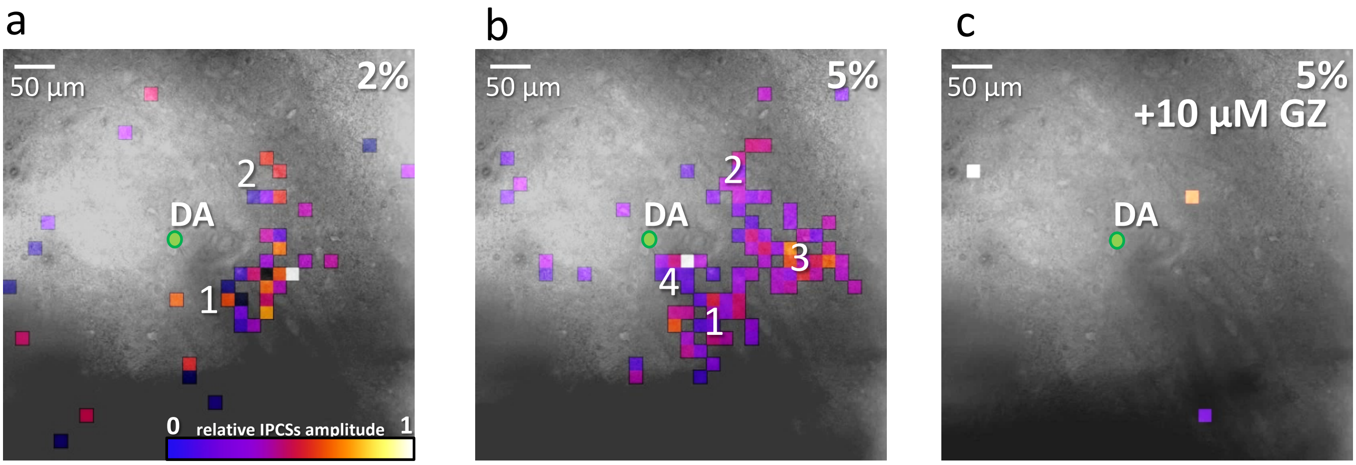


**Figure S8. Representative example of the optical mapping. a – b.** Input maps from 2 (at 2%) and 4 (at 5%) VTA Sst cells to a single DA neuron (green circled dot in the center). **c.** The input map to the same DA neuron, after applying 5% laser stimulation, in the presence of 10 µM Gabazine (GZ), which abolished all evoked IPSCs, confirming GABAergic nature of the inhibitory inputs from the Sst cells.

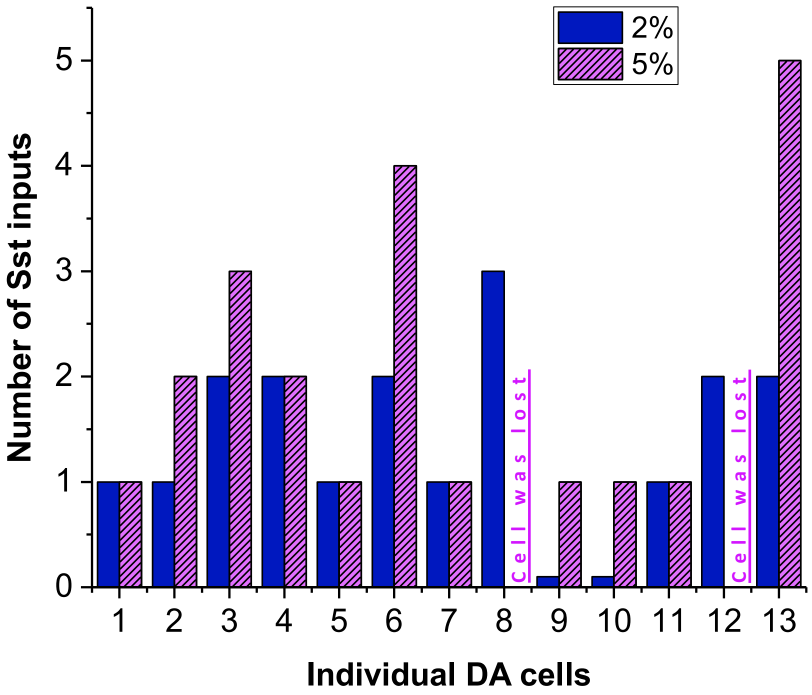


**Figure S9. Number of inputs to DA cells at 2 and 5% laser powers.** Only positive optical maps are shown. Minimum number of inputs for both light intensities was 1. DA cell #13 had the greatest number of inputs, receiving inhibition from five Sst ADP neurons.

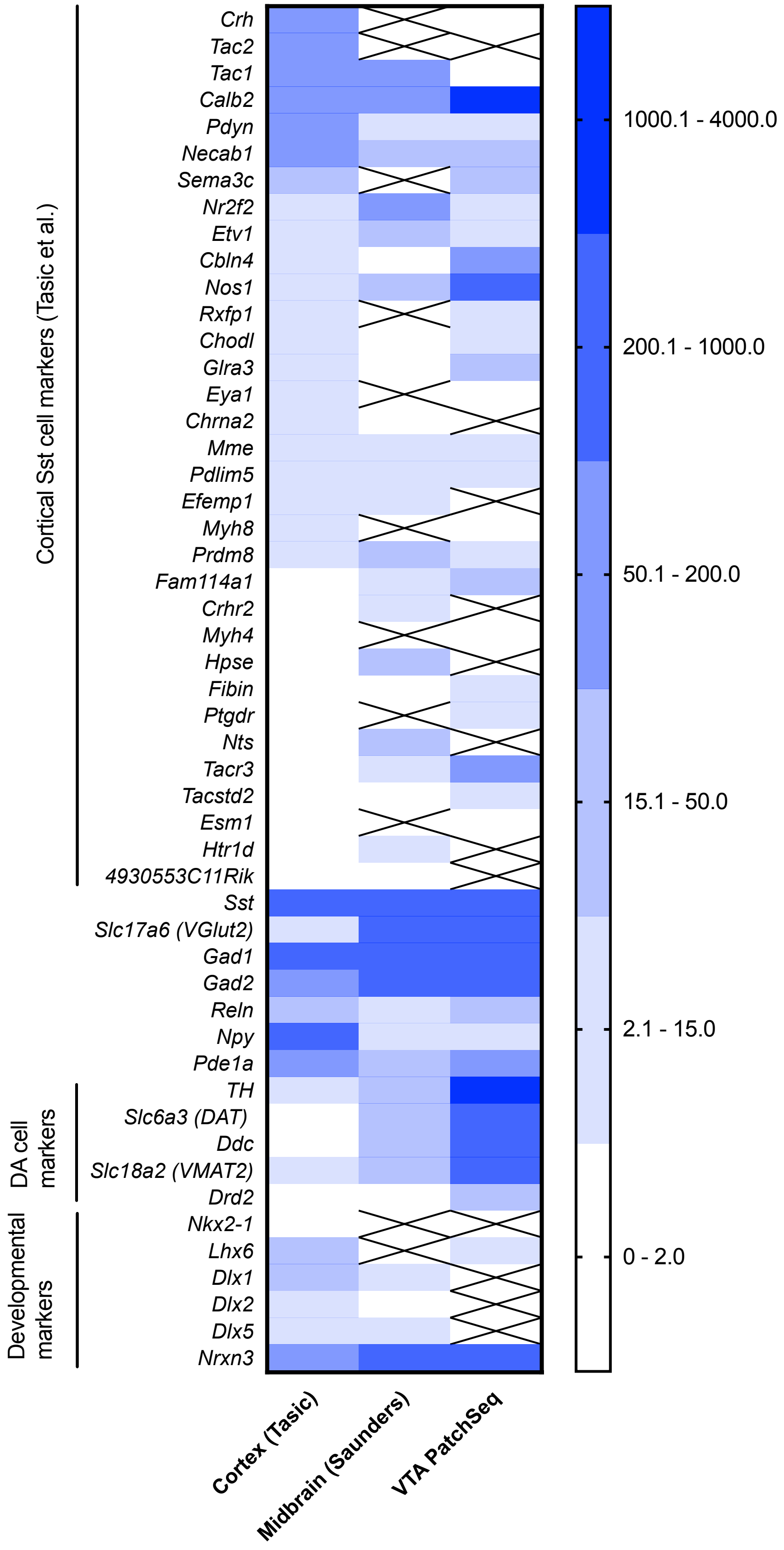


**Figure 8. Heat map of the mean expression of genes of interest in neocortical Sst neurons (Tasic et al. Nature 2018), Sst neurons of the midbrain (Saunders et al. Cell 2018) and in the PatchSeq cells of the present study**. The three sets were normalized by setting the Sst expression to 1000. Crossed out means no expression in the dataset.
